# Hepatocyte androgen receptor in females mediates androgen-induced hepatocellular glucose mishandling and systemic insulin resistance

**DOI:** 10.1101/2021.06.09.447759

**Authors:** Stanley Andrisse, Mingxiao Feng, Zhiqiang Wang, Olubusayo Awe, Lexiang Yu, Haiying Zhang, Sheng Bi, Hongbing Wang, Linhao Li, Serene Joseph, Nicola Heller, Franck Mauvais-Jarvis, Guang William Wong, James Segars, Andrew Wolfe, Sara Divall, Rexford Ahima, Sheng Wu

**Author notes:** Correspondence and name of person to whom reprint requests should be addressed: Sheng Wu, 3500 North Broad Street / MERB 456, Philadelphia, PA, 19140, 215-7073704.

## Abstract

Androgen excess is one of the most common endocrine disorders of reproductive-aged women, affecting up to 20% of this population. Women with elevated androgens often exhibit hyperinsulinemia and insulin resistance. The mechanisms of how elevated androgens affect metabolic function are not clear. Hyperandrogenemia in a dihydrotestosterone (DHT)-treated female mouse model induces whole body insulin resistance possibly through activation of the hepatic androgen receptor (AR). We investigated the role of hepatocyte AR in hyperandrogenemia-induced metabolic dysfunction by using several approaches to delete hepatic AR via animal-, cell-, and clinical-based methodologies. We conditionally disrupted hepatocyte AR in female mice developmentally (LivARKO) or acutely by tail vein injection of an adeno-associated virus with a liver-specific promoter for Cre expression in AR^fl/fl^ mice (adLivARKO). We observed normal metabolic function in littermate female Control (AR^fl/fl^) and LivARKO (AR^fl/fl^; Cre^+/-^) mice. Following chronic DHT treatment, female Control mice treated with DHT (Con-DHT) developed impaired glucose tolerance, pyruvate tolerance, and insulin tolerance, not observed in LivARKO mice treated with DHT (LivARKO-DHT). Further, during an euglycemic hyperinsulinemic clamp, the glucose infusion rate was improved in LivARKO-DHT mice compared to Con-DHT mice. Liver from LivARKO, and primary hepatocytes derived from LivARKO, and adLivARKO mice were protected from DHT-induced insulin resistance and increased gluconeogenesis. These data support a paradigm in which elevated androgens in females disrupt metabolic function via hepatic AR and insulin sensitivity was restored by deletion of hepatic AR.

## 1. Introduction

Androgen excess is one of the most common endocrine disorders of reproductive-aged women, affecting up to 20% of this population[1]. Disorders of hyperandrogenemia in women, such as polycystic ovary syndrome (PCOS) and congenital adrenal hyperplasia are associated with metabolic dysfunction and infertility[2–4]. Women with androgen excess are at a much higher risk than weight-matched women without androgen excess of developing dysglycemia, and the majority (70%) manifest insulin resistance whether lean or obese[5], with 10% developing type-2 diabetes mellitus (T2D)[6].

Similar to women, animal models of female hyperandrogenemia (HA) display metabolic and reproductive dysfunction[2,7–13]. Despite the importance of the liver in systemic insulin action, little is known about the role of hepatocytes in androgen-induced insulin resistance *in vivo* in females. Using a model of low-dose dihydrotestosterone (DHT) to achieve serum levels of DHT 2-fold higher than controls, similar to those of women with PCOS or corrected late-onset congenital adrenal hyperplasia (CAH)[14,15], we showed that androgens modulate liver glucose metabolism to mediate hyperandrogenemia-induced metabolic dysfunction in female mice [16]. This 2XDHT model reflects the androgen level elevation encountered in female human disease and is distinct from other animal models of female hyperandrogenemia[9,17–19] in that body composition (lean mass, fat mass, total weight), serum levels of estradiol, testosterone, and luteinizing hormone (LH) [16,20] during two to three months treatment are similar between treated and untreated mice. This model allows us to parse out the weight independent effects of hyperandrogenemia on glucose metabolism.

While hepatic AR is not required for metabolic function in female mice with normal androgen levels[21], the role of hepatic AR in hyperandrogenic environments has not been explored. The liver plays a major role in determining systemic insulin resistance and metabolism[22] and its function is altered in women[23] and mice[24] with hyperandrogenemia. Here, we investigate the direct role of hepatocyte AR in hyperandrogenemia-induced metabolic dysfunction by using several approaches to delete hepatic AR via animal-, cell-, and clinical-based methodologies. We hypothesize that hepatic AR mediates hyperandrogenemia-induced metabolic abnormalities in female mice and that its deletion will ameliorate hepatic and peripheral metabolic dysfunction. This knowledge may contribute to development of novel and targeted therapeutics to limit the metabolic sequelae of hyperandrogenism experienced by females.

## 2. Materials and Methods

### 2.1. Generation of liver specific AR knockout females

AR^fl^ mice (exon 2) on a C57BL/6 background were generated by Karel De Gendt [25]. Frozen embryos obtained from the European Mutant Mouse Archive (Rome, Italy), were re-derived by the Johns Hopkins Transgenic Core. To produce developmental hepatic AR knockout mice (LivARKO, AR^fl/fl^; albumin (Alb)-Cre^+/-^), we crossed an exon 2-floxed AR [25,26] female (AR^fl/fl^; Cre^-/-^) mouse, with a male albumin-Cre^+/-^ mouse, (strain number: B6N.Cg-Speer6-ps1Tg(Alb-Cre)21Mgn/J) purchased from Jackson Laboratories, Bar Harbor, ME. The Alb-Cre mouse produces liver-specific expression of Cre driven by the albumin promoter in a C57/BL6 background. Litter mates (AR^fl/fl^; Alb-Cre^-/-^) were referred to as Control (Con) mice. Alb-Cre mice are not known to display any metabolic phenotype related to the transgenic allele [27]. Once female mice reached 2-months old, 4 mm-DHT or vehicle pellets were inserted to the mice under the skin [16,20,28–31]. To acutely knockout AR, we used an adeno-associated virus expressing the Cre bacteriophage recombinase, AAV8.TBG.PI.-Cre.rBG. The thyroid binding globulin (TBG) promoter is a gene specifically expressed in the liver [32]. One-hundred microliters of AAV8.TBG.PI.-Cre.rBG [32] (referred to as adLivARKO mice) or pAAV.TBG.PI.Null.bGH (referred to as adCon mice) were injected into AR^fl/fl^ female mice. The liver has been proven as the only targeted tissue by this vector [32]. The two-month-old female mice were inserted with 4 mm-DHT or vehicle pellets simultaneously with tail vein virus injections. Metabolic tests were performed between 1-3-months post insertion. At 2-3-months post insertion, tissues were extracted for further analysis as described below. All mice were housed in the Johns Hopkins University mouse facility, and all experiments were conducted under a protocol approved by the Johns Hopkins Animal Care and Use Committee.

### 2.2. Genotyping and DNA extraction

Genomic DNA was isolated from ear as previously described [16]. Primers used for AR^flox^ genotyping were AR-R AGCCTGTATACTCAGTTGGGG [25], AR-F AATGCATCACATTAAGTTGATACC, Alb-Cre Mutant were Cre-F (CGACCAAGTGACAGCAATGCT), Cre-R (GGTGCTAACCAGCGTTTTCGT). PCR products were resolved on a 1.5 % agarose gel, and DNA bands were visualized by ethidium bromide staining.

### 2.3. Metabolic Testing: GTT, ITT, PTT and GSIS

Mice metabolic testing was performed as described previously [16]. Briefly, for glucose tolerance tests (GTT) and glucose stimulated insulin secretion (GSIS), mice were fasted overnight (16 h) and received intraperitoneal (*i.p*.) injections of 2 g/kg body weight (BW) glucose. For insulin tolerance tests, mice were fasted for 7 hours ad received *i.p*. injections of 0.3 units/kg BW insulin. For pyruvate tolerance tests (PTT), mice were fasted for 12 hours and received *i.p*. injections of 1 g/kg BW pyruvate (PTT). For the GSIS, blood samples were obtained at time zero and at 30 minutes. Serum insulin and c-peptide levels were measured by insulin or C-peptide ELISA respectively following the manufacture’s protocol (Sigma Aldrich, St. Louis, MO, USA). Mouse metabolic testing and fasting methods followed established protocols [16,33,34].

### 2.4. Glucose infusion rate

The euglycemic-hyperinsulinemic glucose clamp is the gold-standard for glucose homeostasis testing (54). The procedure was performed as previously described [33,35,36]. Euglycemia (100-120 mg/dL) was maintained by a variable continuous infusion of a 25% glucose solution adjusted every 10 min. The steady-state glucose infusion rate (GIR) was determined in the last 30 minutes of the 120-minute study.

### 2.5. Metabolic hormones measurement

Six to eight weeks after DHT or empty pellet insertion, serum was obtained from mice fasted for 7 hour and analyzed for insulin, leptin, adiponectin, IL-6 and TNFα using Milliplex Map Mouse Serum Adipokine Panel (Millipore catalogue # MADKMAG-71K) on a Luminex 200IS platform.

### 2.6. Echo MRI: Body Composition Analysis

Eight to ten weeks post pellet insertion, non-fasted Con and LivARKO mice were placed in an Echo MRI apparatus (EchoMRI LLC, Houston, TX) without being restrained or anesthetized to determine body composition, measuring whole body fat, lean, free water, and total water masses in live mice.

### 2.7. qRT-PCR

RNA isolation was performed on collected tissues from WT-Con, WT-DHT, LivARKO-Con, and LivARKO-DHT mice using Trizol (BioRad). Reverse transcription was performed using iScript cDNA synthesis kit (BioRad) and real time quantitative polymerase chain reaction (qRT-PCR) was performed using iQ SYBR Green reagent (BioRad) and an iCycler iQ5 Q-PCR machine (BioRad). Primers were used as previously reported [16].

### 2.8. Western Blot

As described previously [16], liver tissues from mice fasted for 16 h were extracted, quantified via BCA assay, and equivalent quantities of protein were separated via SDS-PAGE (Thermo Scientific, Waltham, MA) and transferred to nitrocellulose membranes. The primary antibodies used were p-AKT, AKT, p-Foxo1 S256, Foxo1, and actin from Cell Signaling Technology (Danvers, Massachusetts) and PI3K-p85, AR (N20) from Santa Cruz Biotechnology (Dallas, Texas), Glucose 6 phosphatase catalytic subunit (G6PC) from Novus Biotechnologicals (Centennial, CO), and phosphoenolpyruvate carboxykinase (PEPCK) from Abcam (Cambridge, MA) all 1:1000 dilutions. The blots were then placed in secondary antibodies (goat anti mouse or goat anti-rabbit, BioRad), and detected using enhanced chemiluminescence (Perkin Elmer Life Sciences, Boston, MA) or Odyssey CLx Imaging System (LI-COR Biosciences, Lincoln, NE).

### 2.9. Immunoprecipitation (IP) and PI3K Activity Assay

Lysates from liver tissues were placed in solution with AR (Santa Cruz) antibodies overnight at 4°C. Protein G sepharose beads (Sigma) were then placed in the mixture for 4 hours at 4°C. Samples were immunoprecipitated as done previously [16]. Immunoprecipitates from pY (Cell Signaling Technology) were subjected to a PI3K activity assay (catalog no. 17-493; Millipore), as detailed in the manufacturer’s protocol manual.

### 2.10. Glucose Uptake Assay

Glucose uptake assays were performed as described previously [28,37] with slight alterations. Mice fasted overnight were euthanized and the tissues were harvested, incubated in Hepes buffered saline (HBS) (plus 32 mM mannitol, 8 mM glucose, and 0.1% radio-immunoassay grade bovine serum albumin (BSA)) for one hour. Then, the glucose uptake assays were performed by incubating the tissues in 12 nM insulin for 20 minutes at 30°C prior to incubation in 2DG transport media (HBS with 4 mM 2DG and 2 μCi/ml 3H-2DG) for 10 minutes at 30°C. The tissues were then washed twice with a 4 mM 2DG rinse (HBS with no radiolabel) for 5 minutes on ice (to remove extracellular 3H-2DG) and stored at −80°C for further analysis. Lysates were counted on a LS6500 liquid scintillation counter (Beckman Coulter, Fullerton, CA) to determine disintegrations per minute (dpm). The DPM were normalized to total protein content.

### 2.11. Female Mouse Primary Hepatocytes

Primary hepatocytes were isolated from livers of 8-10 weeks old Control and LivARKO female mice without DHT treatment. The reagents and materials used for female mouse hepatocyte isolation and culture were described previously [32]. Isolated primary hepatocytes were plated at 0.8 × 10^6^ cells per well of six-well dish in Williams E supplemented with 10% FBS (Gibco). Four to six hours after plating, Fresh media with or without 1 nM DHT was subsequently added for 24 h. Serum starvation was performed for the final 3 h of DHT treatment. Media with or without 100 nM insulin were added for the final half hour of DHT treatment. BioRad cell lysis was used to harvest the cells and the lysate was subsequently processed for western blotting.

### 2.12. Female Human Primary Hepatocytes

University of Maryland Institutional Review Board approved the obtainment, use and distribution of human tissues at the University of Maryland Brain and Tissue Bank. After an approved application for use of the hepatocytes, human hepatocytes (three females, detailed in Suppl. Table 1) were isolated from resection of liver specimens from the Brain and Tissue Bank by a modification of the two-step collagenase digestion method [38,39]. The primary hepatocytes from each individual were treated in two to four separate groups. Group 1-DHT: incubated for 24 hours with media containing vehicle, 0.6 nM, 1 nM, or 6 nM DHT and for the last 4 h of the 24 h incubation, medium was replaced by serum free media. Group 2-DHT+flutamide: the same treatments as in Group1, with 10 nM flutamide (AR inhibitor) added at experiment start. Group 3-DHT+insulin: the same treatments as in Group 1, plus 100nM insulin added in the last 15 min of the 24h. Group 4-DHT+flutamide+insulin: the same treatments as in Group 2, plus 100nM insulin inthe last 30 min of the 24h incubation period. At the end of the experiment, all cells of each well were harvested using cell lysis buffer containing protease and phosphatase inhibitors, and stored at −80°C until further analysis. 0.6-1 nM DHT is equivalent to roughly 2-3 fold of women with PCOS compared that of unaffected women as calculated in [14]. Thus, 6 nM is representative of a supra-pathophysiological dose, and 1nM would be pathophysiological.

### 2.13. Statistical analysis

Statistical analyses used were described in each individual figure legend. Some data were analyzed by student t-tests. Some data were assessed by 2-way ANOVA with main effects of DHT treatment and genotype assessed, or in some experiments DHT exposure and insulin treatment effects. All analyses were performed using Prism software (GraphPad, Inc.). All results were expressed as means ± SEM. A value of p<0.05 was defined as statistically significant.

## 3. Results

### 3.1. Deletion of AR in liver by conditional or adeno-associated virus knockout

To examine the role of hepatocyte AR in the pathophysiology of androgen excess in female mice, we generated a hepatocyte-specific AR knockout mouse. To validate the absence of AR in liver, we measured AR mRNA expression via qRT-PCR and protein expression by western blot. The AR mRNA levels in livers of mice with developmental deletion of hepatic AR (AR^fl/fl^; Cre^+/-^ or LivARKO-veh; Veh refers to vehicle-treated mice) were significantly reduced, yet were not altered in skeletal muscles and gonadal adipose tissues compared with control littermates (AR^fl/fl^ or Con-veh; Figure 1 A-C). AR protein levels of LivARKO were barely detectable in liver (Suppl. Figure 1A-B) compared to Con-veh mice. The AR protein levels were not altered in skeletal muscles or gonadal adipose tissues (Suppl. Figure 1 A, C-D). Furthermore, the liver AR mRNA levels of adult AR^fl/fl^ mice injected with a liver-specific Cre driven adeno-associated virus (AAV8) (adLivARKO or adLivARKO-veh mice) were significantly lower compared with liver AR mRNA levels of adult AR^fl/fl^ mice injected with AAV8-GFP (ad-Con or adCon-veh mice). adLivARKO AR mRNA levels were not altered in skeletal muscles and WAT compared to ad-Con mice (Figure 1 D-F). Mice with genotyping of AR^fl/fl^; AR^wt/wt^ and Alb-Cre^+/-^ do not exhibit different metabolic phenotype, thus many studies have AR^fl/fl^ mice as control (wild type) comparisons[40]. Thus we also used AR^fl/fl^ littermates as controls in this study. Figure 1G details the naming convention we used for the ARKO models in this study.

**Figure 1.**
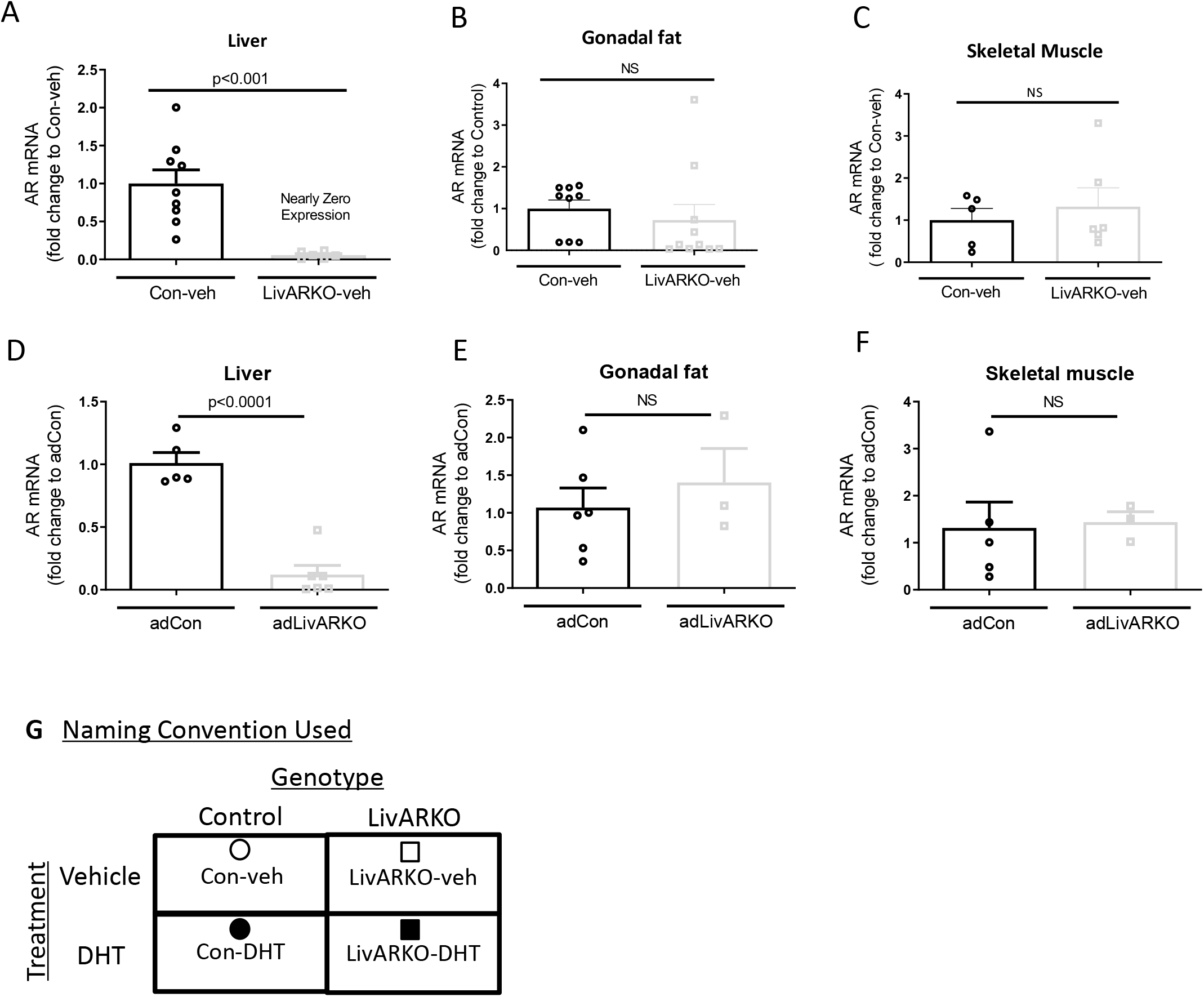
Androgen receptor (AR) is specifically deleted in liver. (A-C) AR mRNA levels of LivARKO compared to control littermates, measured by qRT-PCR, in livers (A), gonadal adipose tissues (WAT, B) and skeletal muscles (C). (D-F) AR mRNA levels in livers (D), WAT (E) and skeletal muscles (F) of adLivARKO mice compared to GFP mice. Two-tailed Students t-tests were applied. P-values were stated in the graphs. Values were mean±S.E.M. (G) Naming and symbol convention used to describe and represent the animals in this study.

- Con-veh = Control mice implanted with an empty pellet represented by an empty circle in the data figures.
- Con-DHT = Control mice implanted with a 4 mm DHT pellet represented by a filled circle.
- LivARKO-veh = mice with liver AR knocked out (KO) conventionally and implanted with an empty pellet represented by an open square.
- LivARKO-DHT = mice with liver ARKO conventionally and implanted with a 4 mm DHT pellet represented by a filled square.

### 3.2. Deletion of AR in liver prevents DHT-induced insulin resistance and impaired glucose homeostasis

As previously reported[21], in basal conditions without DHT and fed normal chow, LivARKO-veh female mice show similar metabolic function as Con-veh mice including normal glucose, insulin, and pyruvate tolerance tests (Figure 2).

**Figure 2:**
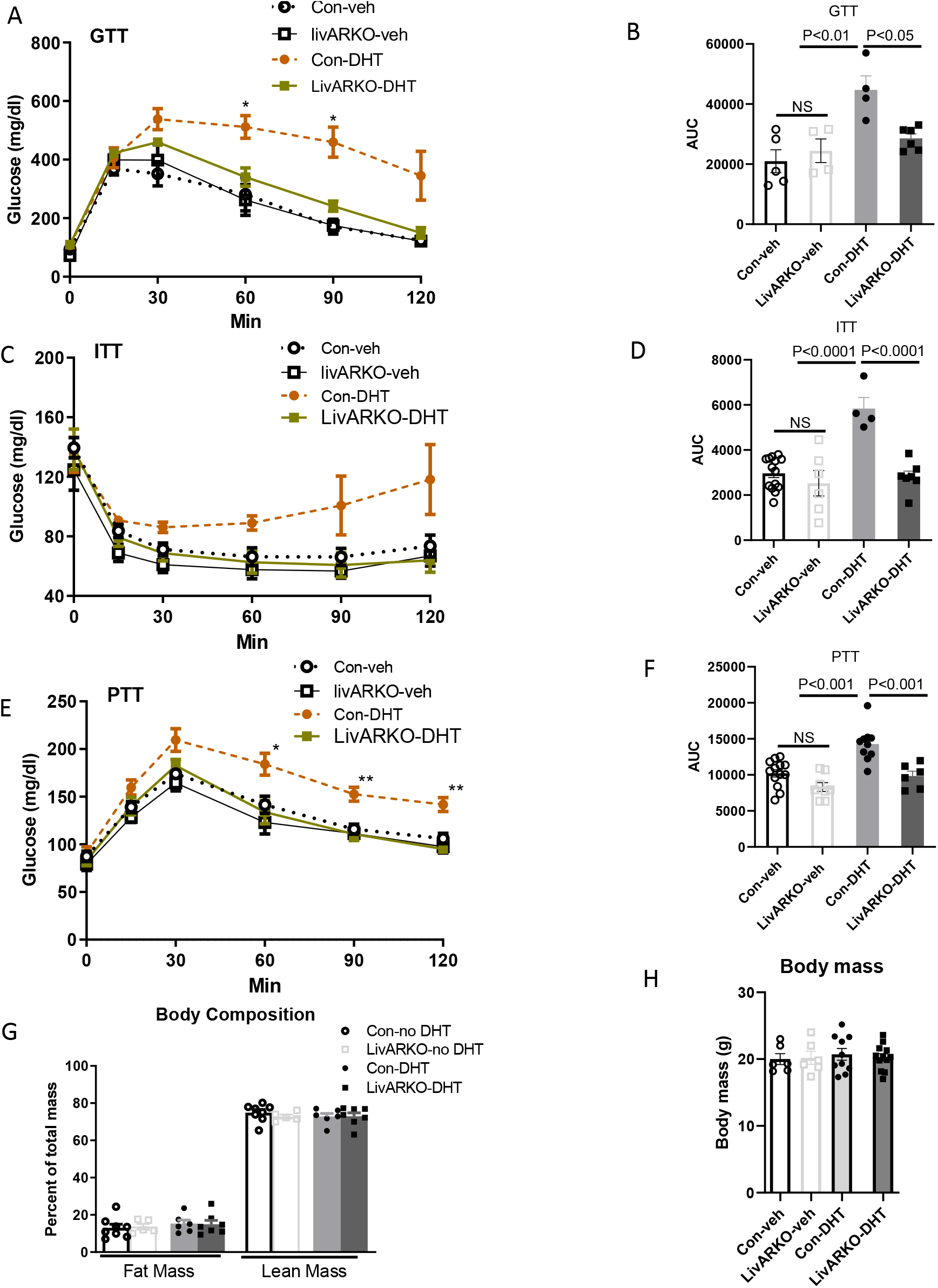
Conditional KO of Liver AR prevents DHT-induced impaired glucose homeostasis. Control and LivARKO mice, with (6-8 weeks after DHT insertion) and without DHT treatment were subjected to (A-B) glucose tolerance test (GTT) (N = 4-6/group), (E-F) insulin tolerance test (ITT) (N = 4-14 /group), or (I-J) Pyruvate tolerance test (PTT) (N = 6-13/group). Con-veh and LivARKO-veh (groups without DHT) are shown on the left. Con-DHT and LivARKO-DHT (groups with DHT) are on the right. The area under the curve (AUC) was determined for the (C-D) GTT, (G-H) ITT, and (K-L) PTT. Statistical analysis was performed for the line graph at the same time point between groups and *=P<0.05, **=P<0.01. (M-N) Fat mass and lean mass were determined via echo magnetic resonance imaging 8 weeks after insertion. (O-P) body weight among groups was recorded (N= 5-10/group). Data were analyzed using two-tailed student’s t-tests. Values are means ±S.E.M.

To determine the effect of hyperandrogenemia on female LivARKO mice, we implanted the mice with slow releasing DHT pellets which were replaced every four weeks. Serum DHT levels were around two-fold higher in mice with DHT pellets compared to mice with empty pellets as previously reported (Suppl. Figure 2A). In the presence of hyperandrogenemia, LivARKO-DHT mice exhibited improved glucose tolerance, insulin sensitivity, and pyruvate tolerance compared to Con-DHT mice (Figure 2). Lean mass, fat mass and body weight (Figure 2G and H) were not different among groups.

LivARKO-veh mice had similar insulin concentrations as Con-veh mice at basal (0 min) and at 30 min post glucose stimulation (Figure 3A); Insulin and c-peptide levels in LivARKO-DHT mice were significantly lower than those of Con-DHT mice (Figure 3B and Suppl. Figure 2B). Improved glucose metabolism was further confirmed by the improved glucose infusion rate and reduced hepatic glucose production (HPG) during euglycemic hyperinsulinemic clamp (GIR, Suppl. Figure 2C-D) in LivARKO-DHT compared to Con-DHT mice. HGP is not uncommon to have it negative[41,42] as far as insulin suppression of HGP is significantly greater in LivARKO-DHT compared to Con-DHT. Hormones and metabolites commonly associated with insulin resistance and obesity (i.e., leptin, IL-6, and tumor necrosis factor-α) showed similar patterns among Con-veh, LivARKO-veh, Con-DHT, and LivARKO-DHT mice (Figure 3C-E).

**Figure 3.**
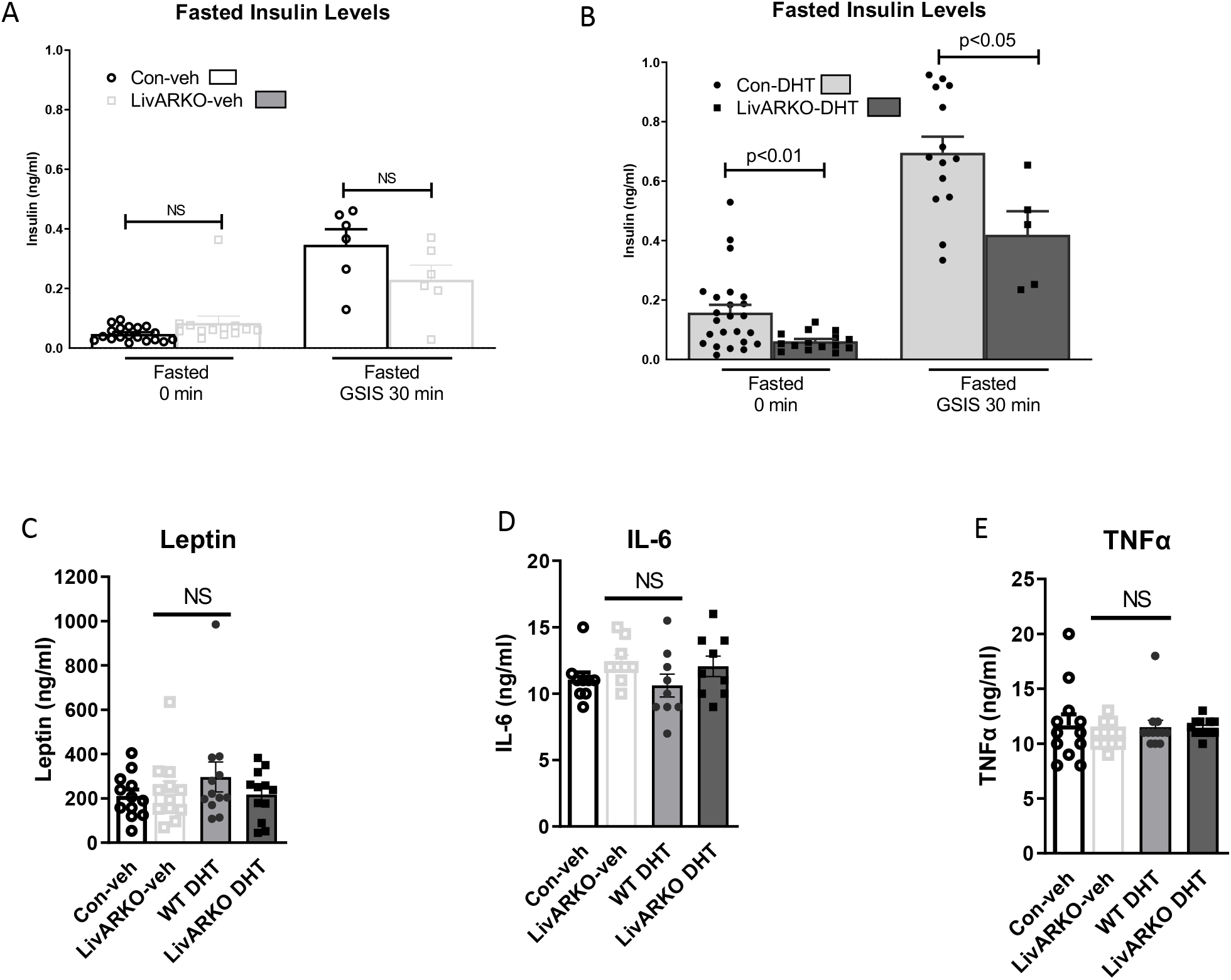
Insulin and hormone levels. Insulin levels were measured from control and LivARKO mice without DHT (A) or at 6-8 weeks after DHT insertion (B) after 16-hour fasted conditions at basal (0 min) and upon GSIS (glucose stimulated insulin secretion at 30min after glucose injection). N = 5-24/group. Hormone levels (C) leptin, (D) IL-6, (E) TNFα, were measured after 7-hour fasted conditions. N=9-12/group. Data were compared by two-tailed student’s t-tests. Values are means ±S.E.M. IL-6 = interleukin 6; TNFα–tumor necrosis factor alpha.

We examined the consequences of adult-onset deletion of hepatic AR by tail vein injection of AAV8-Cre concurrent with DHT insertion in AR^fl/fl^ mice. Adult female mice injected with AAV8-GFP and treated with DHT (adCon-DHT) showed impaired glucose tolerance (IGT) and impaired pyruvate tolerance (IPT), impaired insulin tolerance (IIT) (Figure 4A-F) compared to adCon-veh mice. Concordantly, like LivARKO-DHT mice, adLivARKO-DHT mice exhibited improved GTT and PTT compared to GFP-DHT mice. Additionally, adCon-DHT mice had reduced glucose uptake compared to adCon-veh mice, but this DHT-induced reduction was prevented in the adLivARKO-DHT mice (Figure 4G).

**Figure 4.**
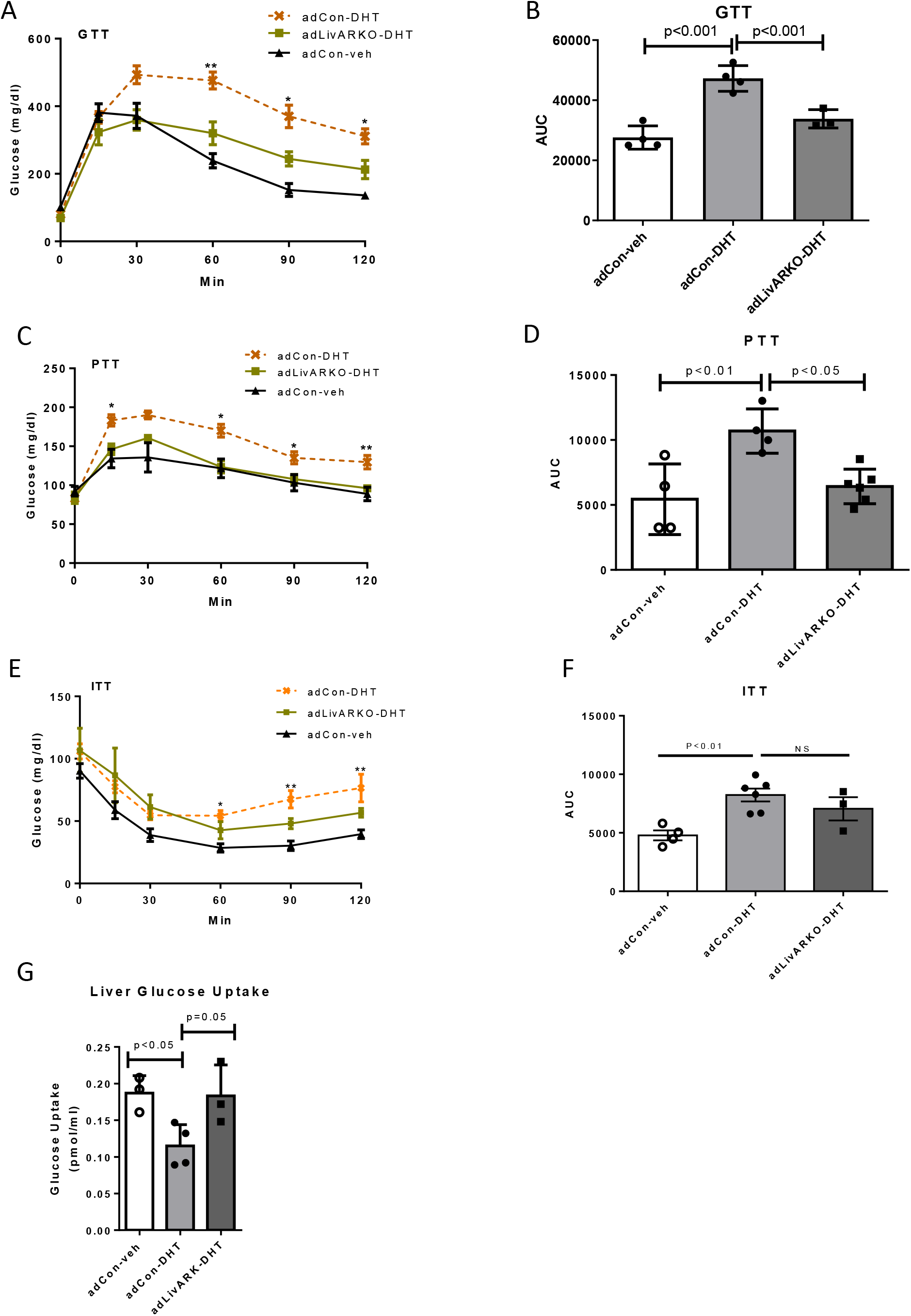
Adult-onset KO of Liver AR prevents DHT-induced impaired glucose homeostasis. DHT and empty pellets were inserted in conjunction with tail vein AAV injections. After 6 weeks post insertion, adCon-veh, adCon-DHT and adLivARKO-DHT were subjected to (A) GTT (N = 3-4/group), (C) PTT (N = 4-6/group), and (D) ITT (N=3-4/group); AUC was determined for the (B) GTT, (D) PTT and (F) ITT. (G). Livers were subjected to ex vivo ^3^H-2-deoxy-glucose glucose transport assays (N=3-4/group). Statistical analysis was performed for the line graph at the same time point using two-way ANOVA followed by Tukey’s multiple tests. *=P<0.05, **=P<0.01, ***=P<0.001 (A,C: adCon-DHT vs adLivARKO-DHT and adCon-veh groups; E adCon-DHT vs adCon-veh groups). Data (B,D,F,G) were analyzed using two-tailed student’s t-tests (adCon-DHT vs adCon-veh or adLivARKO-DHT). Values are mean±S.E.M.

### 3.3. Deletion of hepatic AR ameliorates DHT-induced increase of gluconeogenesis and impaired insulin signaling

AR is a transcription factor, and the impaired glucose production may be due to the dysregulation of gluconeogenic genes. Previously, we observed that female Con-DHT mice had higher expression of gluconeogenic genes (*Pck1*, *G6pc*, and *Foxo1*) and proteins of rate-limiting enzymes (PEPCK and G6Pase) for glucose production [43] in fasted liver compared to Con-Veh female mice[16]. LivARKO-veh mice displayed similar levels of gluconeogenic gene mRNA and PEPCK and G6Pase protein levels as Con-veh mice (Figure 5A, C-E). However, LivARKO-DHT mice showed lower mRNA expression in gluconeogenic genes (*Pck1*, *G6pc, and Foxo1*) (Figure 5B), and protein levels of PEPCK (relative fold change to Con-veh) compared to Con-DHT (Figure 5F-G). We next sought to determine the impact of AR on hepatic insulin signaling in hyperandrogenemia mice. Similar to previous studies[16,28], Con-DHT mice exhibit blunted insulin-stimulated phosphorylation of AKT (Figure 6A-B; noted as insulin resistant) compared to Con-Veh mice. However, this DHT-induced insulin resistance at the level of p-AKT was absent in the LivARKO-DHT mice. We previously observed that AR binds the p85 regulatory subunit of PI3K in liver from Con-DHT mice and impairs insulin signaling at the PI3K level[16]. AR-p85 binding was not observed in livers from LivARKO-DHT mice (Figure 6A, C), and liver PI3K activity was maintained in female LivARKO-DHT mice (Figure 6D). The same phenomenon occurred in the adLivARKO mouse model; DHT disruption of hepatic insulin signaling as indicated by lower insulin stimulated p-AKT was prevented in adLivARKO-DHT mice (Suppl. Figure 4A-B).

**Figure 5.**
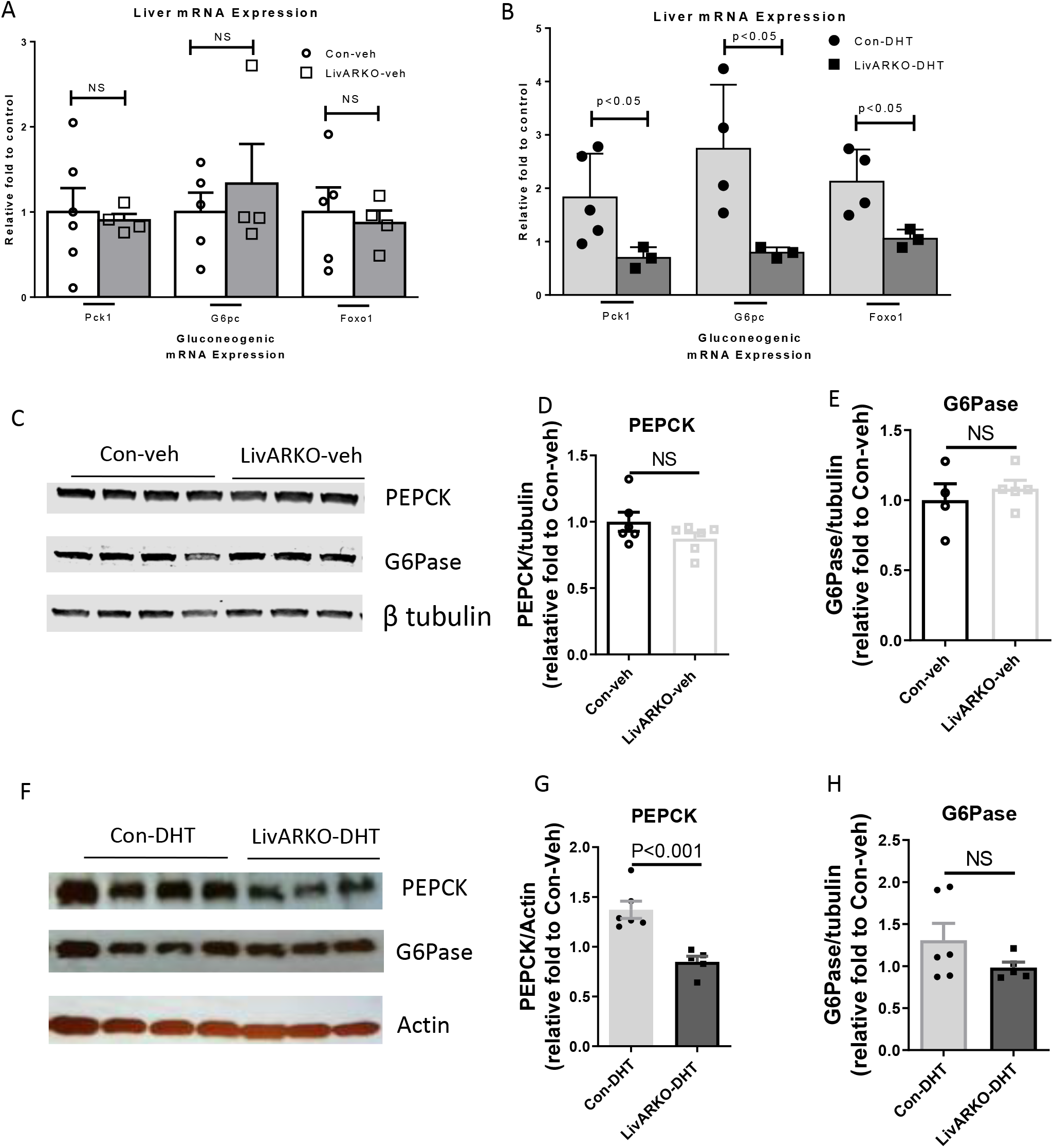
Conditional KO of Liver AR prevents DHT-induced increased expression of gluconeogenic mRNA and proteins. At 10-12 weeks post insertion of pellet, livers of fasted control and LivARKO, with and without DHT mice were taken and processed. (A-B) qRT-PCR analysis to determine mRNA expression of *Pck1*, *G6pc*, *Foxo1* was performed, where (A) shows Con-veh vs. LivARKO-veh and (B) shows Con-DHT vs. LivARKO-DHT. Western blot analysis probing for antibodies against PEPCK and G6Pase and graphical representations of the densitometry, where (C-E) show Con-veh vs. LivARKO-veh and (F-H) show Con-DHT vs. LivARKO-DHT (data are relative fold change to Con-veh). Two-tailed student’s t-tests were used to analyze the data. Values are means ±S.E.M. Pck1 = PEPCK, phosphoenolpyruvate carboxykinase; G6pc, glucose 6 phosphatase catalytic subunit; Foxo1, Forkhead box protein O1.

**Figure 6.**
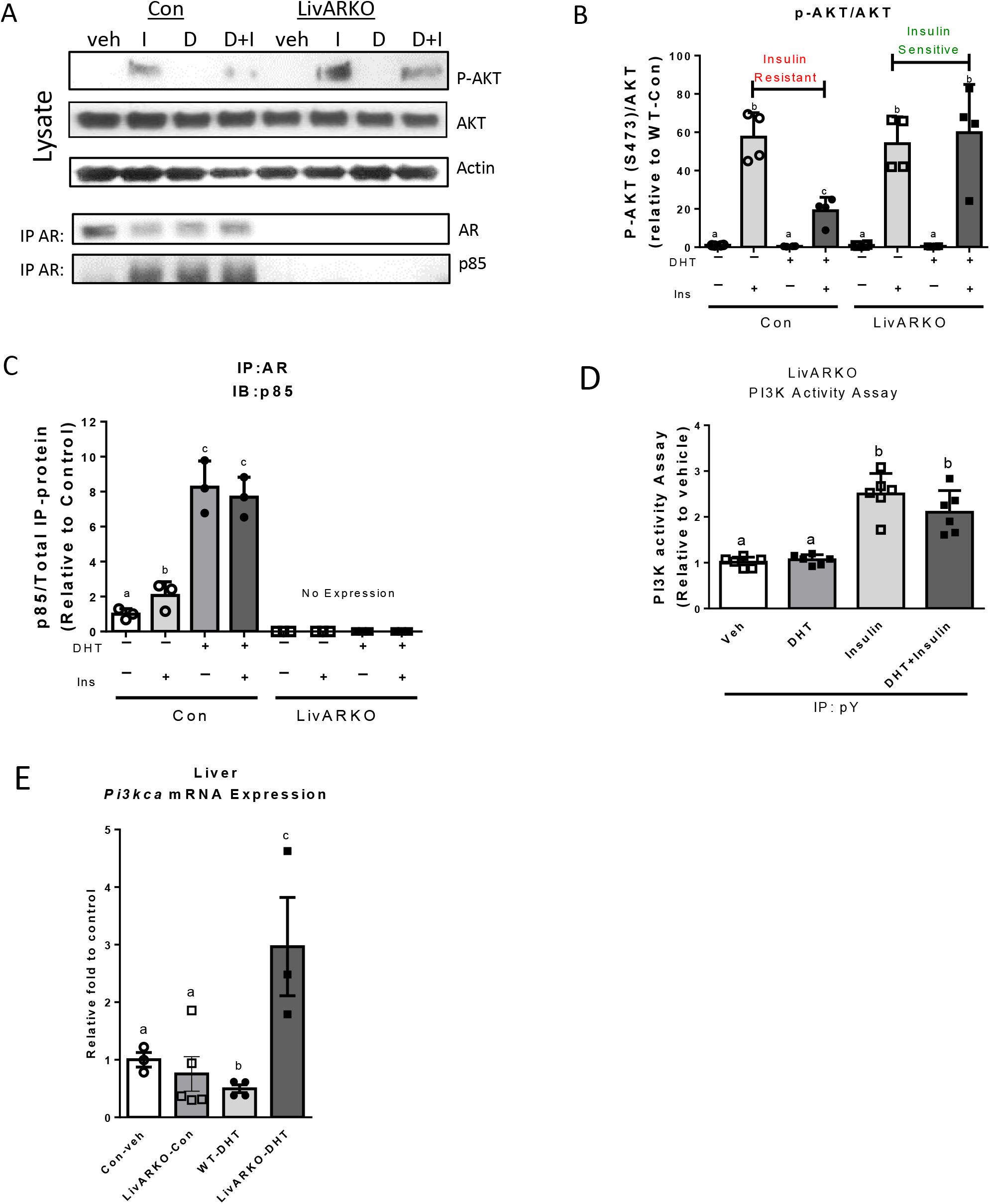
Conditional KO of Liver AR prevents DHT-induced impaired insulin signaling in liver. After 10 weeks post insertion, control and LivARKO mice with or without DHT were fasted for 16 hours and then injected with saline or 0.5 U/kg insulin. After 10 minutes injection, liver samples were collected and subjected to (A) immunoprecipitation and/or western blot analysis. Graphical representations of densitometry for control and LivARKO are shown: (B) p-AKT (S473)/AKT. (C) IP: AR, IB: p85 (N=3-4/group). (D) Liver lysates from LivARKO mice were immunoprecipitated using p-Y antibodies and the IP-pY was subjected to a PI3K activity assay. (N=6/group). A different subset of liver tissues from fasted mice were subjected to (E) qRT-PCR analysis for *Pi3kca* mRNA expression, n = 3-4 per group. Data in this figure were assessed by two-way ANOVA with Tukey’s multiple comparisons tests. Values are means ±S.E.M. Different letters represent statistical difference, P < 0.05. Veh: vehicle; D: DHT; I: insulin; IP: immunoprecipitation; IB: immunoblot.

Although low dose DHT lowered liver mRNA expression of insulin signaling genes (*Pi3kca)* in the fed state[28], livers from LivARKO-DHT mice did not exhibit the DHT-induced decrease in *Pi3kca* (the regulatory p85 subunit of PI3K), compared to livers from WT-DHT (Figure 6E).

### 3.4. Disruption of hepatic AR prevents DHT-induced insulin resistance in primary culture of female mouse and or human hepatocytes

To evaluate if AR directly affects hepatocyte function to induce hepatic insulin resistance rather than indirectly via systemic signaling molecules or tissues, we applied DHT to isolated primary hepatocytes from Control and LivARKO mice *ex vivo*. As observed *in vivo* liver, insulin increased p-AKT and p-Foxo1 in primary hepatocytes. DHT blunted this insulin-stimulated phosphorylation (Figure 7A), thus indicating DHT-induced insulin resistance (marked red in the graphical figure). In primary hepatocytes derived from LivARKO mice, DHT-induced blunting of insulin-stimulated p-AKT and p-Foxo1 was not observed, thus the cells remained insulin-sensitive (marked green in the graphical figure; Figure 7B).

**Figure 7.**
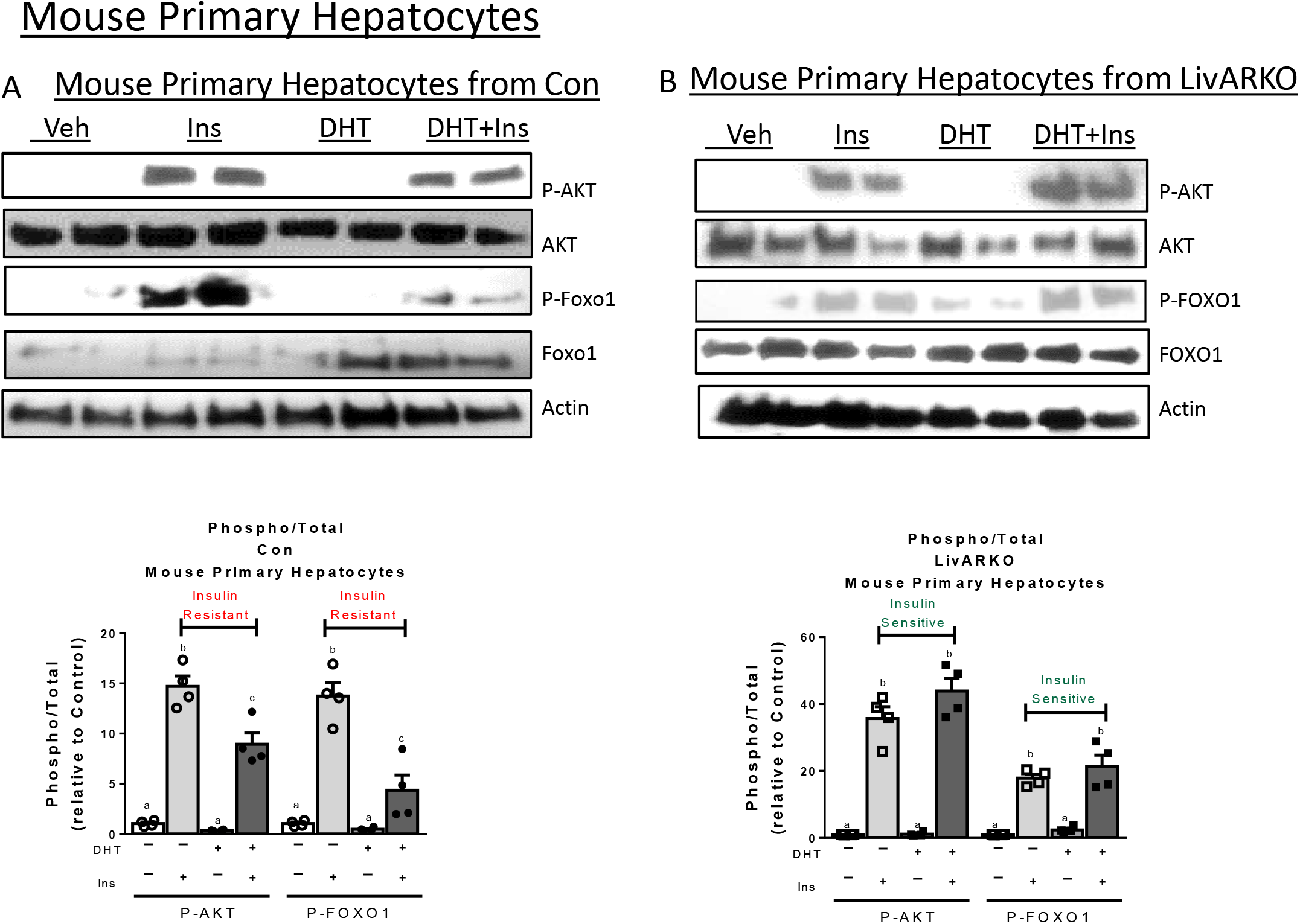
Prevention of DHT-induced insulin resistance was recapitulated in primary hepatocytes from Control and LivARKO female mice. Adult primary hepatocytes were extracted from (A) Control and (B) LivARKO female mice, cultured for 24 hours, and subjected to a cell culture as detailed in the methods. Western blot analysis was performed using antibodies against p-AKT, AKT, p-Foxo1, Foxo1, and Actin. Graphical figures for the densitometry of the blot images are shown below the images. Data were assessed by two-way ANOVA followed by Tukey’s multiple comparisons test. Values are means ±S.E.M. Different letter represents statistical difference, P<0.05.

In human tissue, we examined the effect of DHT in human primary hepatocytes derived from three individual female donors (Suppl. Figure 5). Insulin increased p-AKT and p-FOXO1 in the primary hepatocytes derived from individual 1 (Suppl. Figure 4A-C) and individual 2 (Suppl. Figure 4D-E); and this insulin stimulated increase in p-AKT and p-FOXO1 was significantly reduced in the presence of DHT (Suppl. Figure. 4A-E, marked insulin resistant in the graphical figures). This DHT-induced insulin resistance was prevented in the presence of AR inhibitor, flutamide (Suppl. Figure. 4A-E, marked insulin sensitive in the graphical figure), suggesting that AR-specific androgen induced insulin resistance is occurring in these human hepatocytes as well. Human primary hepatocytes derived from individual 3 female did not exhibit this pattern (Suppl. Figure 4F-G). The n-value for primary, immortalized, and human derived cell culture is a topic of debate[44–47].

## 4. Discussion

Our study shows that hepatocyte AR signaling mediates hyperandrogenemia-induced hepatic insulin resistance in female mice by altering hepatic insulin signaling and phosphorylation cascades to increase gluconeogenesis and systemic glucose intolerance. Hepatocyte AR did not play a role in the developmental formation of insulin signaling in the hepatocyte, as LivARKO-veh female had no metabolic phenotype under physiological androgen levels. In hyperandrogenic states, female mice with developmental or postnatal deletion of AR did not exhibit impaired metabolic function compared to female mice with intact AR.

AR signaling in multiple cells including the theca cell, gonadotroph, adipocyte, and neurons have been found to be involved in the pathophysiology of metabolic and ovarian dysfunction in mouse models of female hyperandrogenemia[20,30,48]. That multiple cell types are involved is not surprising considering the multiorgan dysfunction associated with hyperandrogenemia. In these murine models of hyperandrogenemia, various degrees of hyperandrogenemia are achieved experimentally, which may account for the various degrees of weight gain, body composition, and metabolic dysfunction observed among models. Our low dose DHT model offers an opportunity to investigate metabolic effects of hyperandrogenemia in a lean state as body weight, composition, and systemic markers of insulin resistance are similar among the groups (Figure 2 and 3) during two to three months of DHT treatment.

Low dose DHT blunts insulin action and alters glucose uptake in liver and adipose tissues without affecting insulin action in skeletal muscle [16,28]. The impaired glucose tolerance is not likely due to pancreatic β cell insufficiency since there is higher basal insulin and C-peptide levels in low dose DHT versus vehicle treated mice (Figure 3B and Suppl. Figure 2B). In contrast to our findings, female mice exposed to a higher dose of DHT (4-fold higher than untreated) on a western (high carbohydrate and high fat) diet exhibit pancreatic β cell dysfunction driven by β cell AR and central neuronal insulin resistance with resultant failure of insulin to suppress hepatic glucose production [49,50]. In another study, female mice exposed to higher dose DHT (8-fold increase in serum DHT levels)[51] than our study exhibit weight gain, adipocyte hypertrophy, and hepatic steatosis that is mostly ameliorated by loss of AR (AR floxed on exon 3) in the adipocyte and neuron [48]. Unlike other models of DHT treatment, we do not observe weight gain, increased basal blood glucose levels or hepatic steatosis [16,28]. This may be due to dose of DHT or exposure time to DHT. Combined, these studies suggest that there is differential cellular sensitivity to AR receptor occupation, with the hepatocyte sensitive to a lower amount of AR receptor signaling to invoke glucose mishandling while the beta cell require higher dose of androgen to trigger cellular dysfunction via AR signaling.

These results are in agreement with data from lean women with hyperandrogenism due to PCOS [52]. Dunaif et al., found that baseline hepatic glucose production is much higher, and that hepatic glucose production was less suppressed upon insulin infusion in lean PCOS women compared to lean non-PCOS women, suggesting that hepatic glucose production in women with PCOS is influenced by other factors besides insulin. Insulin levels of PCOS women are typically more than two-fold higher than non PCOS women [52]. Thus, lean PCOS women have greater hepatic glucose production upon fasting even with a higher circulating insulin level. Like what is seen in lean PCOS women, the liver was insulin-resistant in our low dose DHT mouse model. Impaired hepatic insulin signaling and glucose homeostasis contributions to the metabolic phenotype of lean hyperandrogenic females is confirmed with the studies herein, as LivARKO-DHT mice do not experiences DHT-disrupted glucose metabolism.

Androgen excess impairs glucose handling via direct AR actions in the hepatocyte (Figure 2). Hepatic glucose production occurs via glycogenolysis and gluconeogenic enzyme activity [53]. LivARKO mice did not exhibit DHT-induced increase in mRNA and protein expression of the gluconeogenic genes *Pck1* or*G6pc* (Figure 5). Foxo1 is a gluconeogenic transcription factor and reducing Foxo1 expression will dampen gluconeogenesis by reducing *Pck1* and *G6pc*. AR inhibition of insulin-stimulated Foxo1 degradation results in increased Foxo1 protein levels and consequent increased PEPCK and or G6Pase protein levels in liver of control but not LivARKO mice (Figure 5 and [16]). The increased gluconeogenesis in control DHT treated mice was not observed with an increase in higher fasting glucose levels (Figure 2) likely because the insulin levels trended higher in control-DHT treated animals (Figure 3).

AR also directly modulates the hepatocellular insulin signaling cascade. Insulin-induced phosphorylation of AKT is reduced in control mice pretreated with DHT (Figure 6A-B) while p-AKT protein levels are preserved in LivARKO-DHT mice indicating that DHT modulates downstream insulin signaling (Figure 6A-B). Immunoprecipitation of p85 and AR occurred in the presence of DHT with or without insulin and did not occur in LivARKO mice, (Figure 6C), indicating that AR directly binds p85. Insulin induced hepatocyte PI3K activity in LivARKO mice was unaltered by the presence of DHT (Figure 6D) in contrast to control mice [16]. DHT alteration of insulin-induced phosphorylation of AKT and Foxo1 also occur in primary hepatocytes from control but not LivARKO mice (Figure 7) indicating that AR action on insulin signaling is not dependent upon other paracrine or endocrine signaling. Hyperandrogenism also disrupts insulin signaling in female human liver as 2 of 3 human samples displayed differential insulin-induced phosphorylation of AKT and FOXO1 in the presence of DHT. Differences in body mass index (BMI was only 19.5), race, and medical history may account for the different pattern in the 3^rd^ human sample. Our previous studies indicate that AR binds directly to p85 *in vitro*, which in turn may blunt cytosolic signaling activity of PI3K and insulin induced phosphorylation (16); disruption of p85α / p85β binding to p110 blocks docking of p110 to IRS1/2 thus decreases PI3K activity [16,54]. Via modulating insulin induced phosphorylation cascades critical for regulating gluconeogenesis and glycogenolysis, androgen action via the androgen receptor modulates the hepatocyte response to insulin, leading to hepatocellular insulin resistance.

Azziz et al., [1] indicated that the mechanism of insulin resistance in obese PCOS women was increased serine phosphorylation of insulin receptor and IRS1 in myocytes, which inhibits metabolic insulin action. Our DHT mice showed similar body mass and body composition and suggest a different or additional mechanism in the non-obese state involving the interaction of hepatic AR at the level of PI3K leading to hepatocyte insulin resistance. Hyperandrogenic-induced alteration in hepatocyte lipid metabolism may also affect hepatocyte cellular function and insulin signaling to account for some of the results observed. However, we have not noted a difference in fat accumulation as assessed by oil-red-O staining in our low dose DHT model [16] suggesting that hepatic steatosis is not a factor in the hepatocyte insulin resistance. It is possible that with longer DHT treatment at this low dose, hepatic lipid metabolism may be altered further affecting hepatocellular function and glucose handling. Subtle effects on lipogenesis and lipid handling may occur with the doses and duration of DHT used in these experiments but are beyond the scope of the current manuscript. Nuclear steroid receptors may regulate mitochondrial function and thus cellular ATP[55] to affect cellular function and/or enhance gene transcription. We cannot exclude the possibility that the observed effect of AR on gluconeogenesis is in part due to effects on mitochondrial function. Our *in vitro* studies indicate that AR directly binds enzymes in the insulin signaling cascade, a task that would not be affected by mitochondrial function. Hyperandrogenemia-induced inhibition of insulin signaling could also be regulated by other novel pathways such as the intracellular tyrosine kinase C (Src) [56] and extracellular signal-regulated kinase (Erk1/2) [54,57], a focus of future study.

In conclusion, we show that androgen action via hepatocyte AR mediates hepatic insulin resistance and glucose metabolism upon hyperandrogenemia in female mice (Figure 8) and murine and human primary hepatocytes. Hepatic AR could thus be a target to direct therapies specifically for women with hyperandrogenemia-associated altered glucose homeostasis.

**Figure 8.**
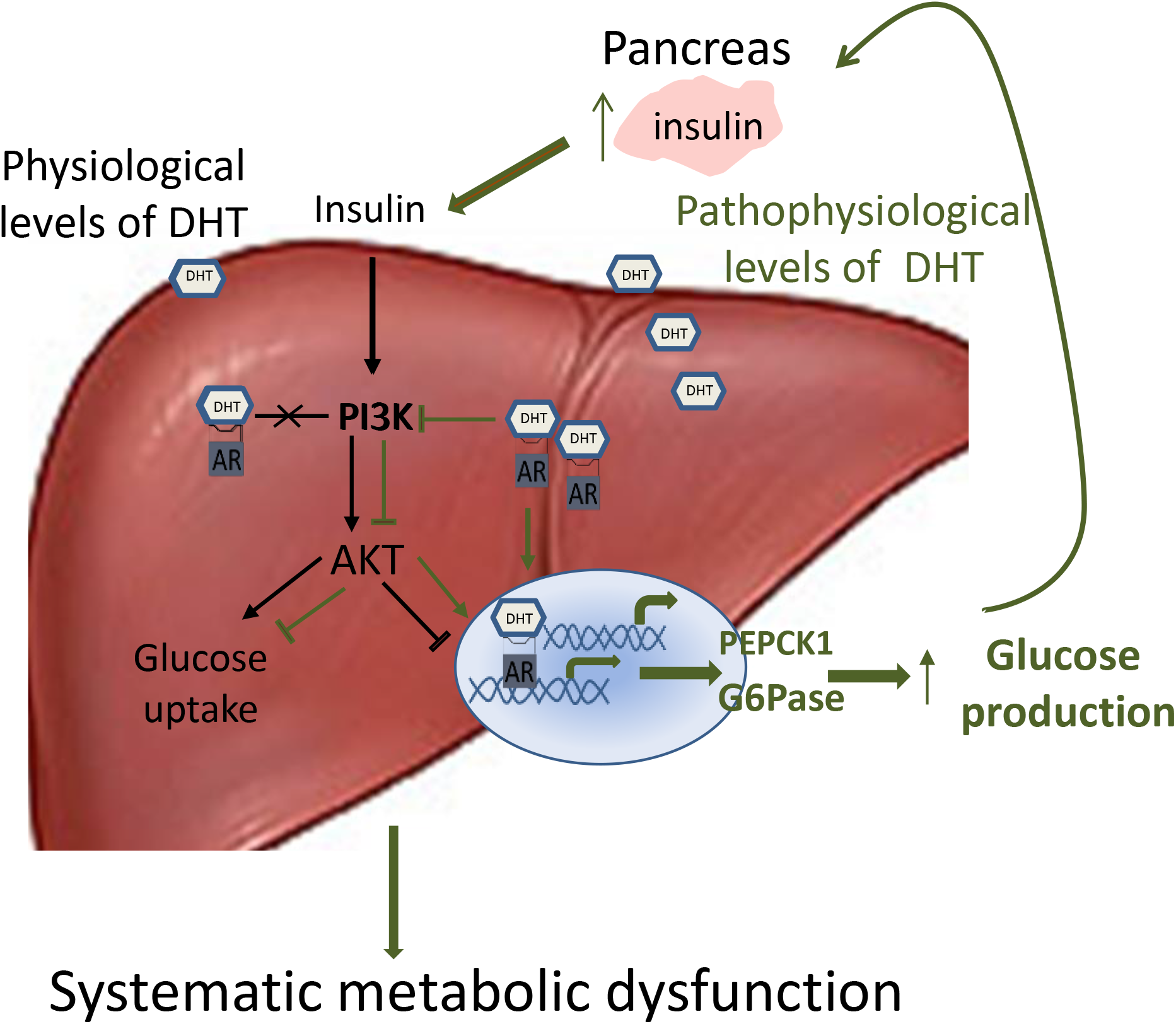
The effects of physiological and pathophysiological levels of androgen in female liver. Under physiological levels of DHT (black line), insulin signaling and gluconeogenesis proceed normally. Under pathophysiological levels of DHT (green line), cytosolic liver AR impairs insulin signaling and nuclear AR directly regulates gluconeogenesis resulting in enhanced gluconeogenesis. This is associated with hyperinsulinemia and systemic insulin resistance.

## Abbreviations

*AR*: *androgen receptor*
PCOS: polycystic ovary syndrome
DHT: dihydrotestosterone
CAH: congenital adrenal hyperplasia
HA: hyperandrogenemia
Con: control
Veh: vehicle
LH: luteinizing hormone
Foxo1: forkhead box protein O1
PEPCK: phosphoenolpyruvate carboxykinase
G6Pase: glucose 6-phosphatase
*I.P*.: intraperitoneal
DPM: determine disintegrations per minute
BMI: body mass index (BMI)
GTT: glucose tolerance test
ITT: insulin tolerance test
PTT: pyruvate tolerance test
GSIS: glucose stimulated insulin secretion
HGP: hepatic glucose production
GIR: glucose infusion rate.

## Author Contributions

SA and SW contributed to the conceptual design, performance of experiments, interpretation, and analysis of data, and writing and editing the manuscript. All other authors contributed to performing some of the experiments, analyzing the corresponding data, and reviewing and editing the manuscript.

## Acknowledgements

This work was supported by the National Institutes of Health (Grants R00-HD068130 04 and R01-HD09551201A1 to S.W.). FMJ was supported by R01 grants DK074970 and DK107444, a Department of Veterans Affairs Merit Review Award (#BX003725) and the Tulane Center of Excellence in Sex-Based Biology & Medicine. NMH was supported by R01 HL124477 and a DOD Award (W81XWH-16-I-0509). SB was supported by DK103710. Technical support was provided by the Integrated Physiology Core of the Baltimore DRTC (P30DK079637). We thank Dr. Jun Luo (Johns Hopkins University School of Medicine) and Dr. William Henry Walker (University of Pittsburg) for offering us the control and DNA binding mutant AR plasmids in this experiment.

## References

[1] Azziz, R., Carmina, E., Chen, Z., Dunaif, A., Laven, J.S., Legro, R.S., et al., 2016. Polycystic ovary syndrome. Nature reviews. Disease primers 2:16057.

[2] Padmanabhan, V., Veiga-Lopez, A., 2013. Sheep models of polycystic ovary syndrome phenotype. Molecular and cellular endocrinology 373:8–20.

[3] Navarro, G., Allard, C., Xu, W., Mauvais-Jarvis, F., 2015. The role of androgens in metabolism, obesity, and diabetes in males and females. Obesity (Silver Spring, Md.) 23:713–719.

[4] Escobar-Morreale, H.F., Alvarez-Blasco, F., Botella-Carretero, J.I., Luque-Ramirez, M., 2014. The striking similarities in the metabolic associations of female androgen excess and male androgen deficiency. Human reproduction (Oxford, England) 29:2083–2091.

[5] Barber, T.M., Dimitriadis, G.K., Andreou, A., Franks, S., 2015. Polycystic ovary syndrome: insight into pathogenesis and a common association with insulin resistance. Clinical medicine (London, England) 15 Suppl 6:s72–6.

[6] Ehrmann, D.A., 2012. Metabolic dysfunction in pcos: Relationship to obstructive sleep apnea. Steroids 77:290–294.

[7] Stener-Victorin, E., Padmanabhan, V., Walters, K.A., Campbell, R.E., Benrick, A., Giacobini, P., et al., 2020. Animal Models to Understand the Etiology and Pathophysiology of Polycystic Ovary Syndrome. Endocrine reviews 41:10.1210/endrev/bnaa010.

[8] Roland, A.V., Nunemaker, C.S., Keller, S.R., Moenter, S.M., 2010. Prenatal androgen exposure programs metabolic dysfunction in female mice. The Journal of endocrinology 207:213–223.

[9] van Houten, E.L., Kramer, P., McLuskey, A., Karels, B., Themmen, A.P., Visser, J.A., 2012. Reproductive and metabolic phenotype of a mouse model of PCOS. Endocrinology 153:2861–2869.

[10] Walters, K.A., Allan, C.M., Handelsman, D.J., 2012. Rodent models for human polycystic ovary syndrome. Biology of reproduction 86:149, 1–12.

[11] Abbott, D.H., Bruns, C.R., Barnett, D.K., Dunaif, A., Goodfriend, T.L., Dumesic, D.A., et al., 2010. Experimentally induced gestational androgen excess disrupts glucoregulation in rhesus monkey dams and their female offspring. American journal of physiology. Endocrinology and metabolism 299:E741–51.

[12] van Houten, E.L., Visser, J.A., 2014. Mouse models to study polycystic ovary syndrome: a possible link between metabolism and ovarian function?. Reproductive biology 14:32–43.

[13] Cardoso, R.C., Puttabyatappa, M., Padmanabhan, V., 2015. Steroidogenic versus Metabolic Programming of Reproductive Neuroendocrine, Ovarian and Metabolic Dysfunctions. Neuroendocrinology 102:226–237.

[14] Silfen, M.E., Denburg, M.R., Manibo, A.M., Lobo, R.A., Jaffe, R., Ferin, M., et al., 2003. Early endocrine, metabolic, and sonographic characteristics of polycystic ovary syndrome (PCOS): comparison between nonobese and obese adolescents. The Journal of clinical endocrinology and metabolism 88:4682–4688.

[15] Ambroziak, U., Kepczynska-Nyk, A., Kurylowicz, A., Malunowicz, E.M., Wojcicka, A., Miskiewicz, P., et al., 2016. The diagnosis of nonclassic congenital adrenal hyperplasia due to 21-hydroxylase deficiency, based on serum basal or post-ACTH stimulation 17-hydroxyprogesterone, can lead to false-positive diagnosis. Clinical endocrinology 84:23–29.

[16] Andrisse, S., Childress, S., Ma, Y., Billings, K., Chen, Y., Xue, P., et al., 2016. Low Dose Dihydrotestosterone Drives Metabolic Dysfunction via Cytosolic and Nuclear Hepatic Androgen Receptor Mechanisms. Endocrinology:en20161553.

[17] Manneras, L., Cajander, S., Holmang, A., Seleskovic, Z., Lystig, T., Lonn, M., et al., 2007. A new rat model exhibiting both ovarian and metabolic characteristics of polycystic ovary syndrome. Endocrinology 148:3781–3791.

[18] Lai, H., Jia, X., Yu, Q., Zhang, C., Qiao, J., Guan, Y., et al., 2014. High-fat diet induces significant metabolic disorders in a mouse model of polycystic ovary syndrome. Biology of reproduction 91:127.

[19] Marino, J.S., Iler, J., Dowling, A.R., Chua, S., Bruning, J.C., Coppari, R., et al., 2012. Adipocyte dysfunction in a mouse model of polycystic ovary syndrome (PCOS): evidence of adipocyte hypertrophy and tissue-specific inflammation. PloS one 7:e48643.

[20] Ma, Y., Andrisse, S., Chen, Y., Childress, S., Xue, P., Wang, Z., et al., 2016. Androgen Receptor in the Ovary Theca Cells Plays a Critical Role in Androgen-Induced Reproductive Dysfunction. Endocrinology: en20161608.

[21] Lin, H.Y., Yu, I.C., Wang, R.S., Chen, Y.T., Liu, N.C., Altuwaijri, S., et al., 2008. Increased hepatic steatosis and insulin resistance in mice lacking hepatic androgen receptor. Hepatology (Baltimore, Md.) 47:1924–1935.

[22] Santoleri, D., Titchenell, P.M., 2019. Resolving the Paradox of Hepatic Insulin Resistance. Cellular and molecular gastroenterology and hepatology 7:447–456.

[23] Vassilatou, E., 2014. Nonalcoholic fatty liver disease and polycystic ovary syndrome. World journal of gastroenterology 20:8351–8363.

[24] Sanchez-Garrido, M.A., Tena-Sempere, M., 2020. Metabolic dysfunction in polycystic ovary syndrome: Pathogenic role of androgen excess and potential therapeutic strategies. Molecular metabolism 35:100937.

[25] De Gendt, K., Swinnen, J.V., Saunders, P.T., Schoonjans, L., Dewerchin, M., Devos, A., et al., 2004. A Sertoli cell-selective knockout of the androgen receptor causes spermatogenic arrest in meiosis. Proceedings of the National Academy of Sciences of the United States of America 101:1327–1332.

[26] Yeh, S., Tsai, M.Y., Xu, Q., Mu, X.M., Lardy, H., Huang, K.E., et al., 2002. Generation and characterization of androgen receptor knockout (ARKO) mice: an in vivo model for the study of androgen functions in selective tissues. Proceedings of the National Academy of Sciences of the United States of America 99:13498–13503.

[27] Postic, C., Dentin, R., Girard, J., 2004. Role of the liver in the control of carbohydrate and lipid homeostasis. Diabetes & metabolism 30:398–408.

[28] Andrisse, S., Billings, K., Xue, P., Wu, S., 2018. Insulin signaling displayed a differential tissue-specific response to low-dose dihydrotestosterone in female mice. American journal of physiology.Endocrinology and metabolism 314:E353–E365.

[29] Wang, Z., Shen, M., Xue, P., DiVall, S.A., Segars, J., Wu, S., 2018. Female Offspring From Chronic Hyperandrogenemic Dams Exhibit Delayed Puberty and Impaired Ovarian Reserve. Endocrinology 159:1242–1252.

[30] Wang, Z., Feng, M., Awe, O., Ma, Y., Shen, M., Xue, P., et al., 2019. Gonadotrope androgen receptor mediates pituitary responsiveness to hormones and androgen-induced subfertility. JCI insight 5:10.1172/jci.insight.127817.

[31] Xue, P., Wang, Z., Fu, X., Wang, J., Punchhi, G., Wolfe, A., et al., 2018. A Hyperandrogenic Mouse Model to Study Polycystic Ovary Syndrome. Journal of visualized experiments: JoVE (140). doi:10.3791/58379.

[32] Qiu, S., Vazquez, J.T., Boulger, E., Liu, H., Xue, P., Hussain, M.A., et al., 2017. Hepatic estrogen receptor alpha is critical for regulation of gluconeogenesis and lipid metabolism in males. Scientific reports 7:1661-017–01937-4.

[33] Ayala, J.E., Samuel, V.T., Morton, G.J., Obici, S., Croniger, C.M., Shulman, G.I., et al., 2010. Standard operating procedures for describing and performing metabolic tests of glucose homeostasis in mice. Disease models & mechanisms 3:525–534.

[34] Jensen, T.L., Kiersgaard, M.K., Sorensen, D.B., Mikkelsen, L.F., 2013. Fasting of mice: a review. Laboratory animals 47:225–240.

[35] Li, L., de La Serre, C.B., Zhang, N., Yang, L., Li, H., Bi, S., 2016. Knockdown of Neuropeptide Y in the Dorsomedial Hypothalamus Promotes Hepatic Insulin Sensitivity in Male Rats. Endocrinology 157:4842–4852.

[36] Lee, B., Qiao, L., Lu, M., Yoo, H.S., Cheung, W., Mak, R., et al., 2014. C/EBPalpha regulates macrophage activation and systemic metabolism. American journal of physiology.Endocrinology and metabolism 306:E1144–54.

[37] Andrisse, S., Patel, G.D., Chen, J.E., Webber, A.M., Spears, L.D., Koehler, R.M., et al., 2013. ATM and GLUT1-S490 phosphorylation regulate GLUT1 mediated transport in skeletal muscle. PloS one 8:e66027.

[38] LeCluyse, E.L., Alexandre, E., Hamilton, G.A., Viollon-Abadie, C., Coon, D.J., Jolley, S., et al., 2005. Isolation and culture of primary human hepatocytes. Methods in molecular biology (Clifton, N.J.) 290:207–229.

[39] Lecluyse, E.L., Alexandre, E., 2010. Isolation and culture of primary hepatocytes from resected human liver tissue. Methods in molecular biology (Clifton, N.J.) 640:57–82.

[40] Chang, C., Yeh, S., Lee, S.O., Chang, T.M., 2013. Androgen receptor (AR) pathophysiological roles in androgen-related diseases in skin, bone/muscle, metabolic syndrome and neuron/immune systems: lessons learned from mice lacking AR in specific cells. Nuclear receptor signaling 11:e001.

[41] Allard, C., Morford, J.J., Xu, B., Salwen, B., Xu, W., Desmoulins, L., et al., 2019. Loss of Nuclear and Membrane Estrogen Receptor-alpha Differentially Impairs Insulin Secretion and Action in Male and Female Mice. Diabetes 68:490–501.

[42] Batista, T.M., Dagdeviren, S., Carroll, S.H., Cai, W., Melnik, V.Y., Noh, H.L., et al., 2020. Arrestin domain-containing 3 (Arrdc3) modulates insulin action and glucose metabolism in liver. Proceedings of the National Academy of Sciences of the United States of America 117:6733–6740.

[43] Jitrapakdee, S., 2012. Transcription factors and coactivators controlling nutrient and hormonal regulation of hepatic gluconeogenesis. The international journal of biochemistry & cell biology 44:33–45.

[44] Lazic, S.E., Clarke-Williams, C.J., Munafo, M.R., 2018. What exactly is ‘N’ in cell culture and animal experiments?. PLoS biology 16:e2005282.

[45] Cumming, G., Fidler, F., Vaux, D.L., 2007. Error bars in experimental biology. The Journal of cell biology 177:7–11.

[46] Geraghty, R.J., Capes-Davis, A., Davis, J.M., Downward, J., Freshney, R.I., Knezevic, I., et al., 2014. Guidelines for the use of cell lines in biomedical research. British journal of cancer 111:1021–1046.

[47] Naegle, K., Gough, N.R., Yaffe, M.B., 2015. Criteria for biological reproducibility: what does “n” mean?. Science signaling 8:fs7.

[48] Cox, M.J., Edwards, M.C., Rodriguez Paris, V., Aflatounian, A., Ledger, W.L., Gilchrist, R.B., et al., 2020. Androgen Action in Adipose Tissue and the Brain are Key Mediators in the Development of PCOS Traits in a Mouse Model. Endocrinology 161:10.1210/endocr/bqaa061.

[49] Xu, W., Morford, J., Mauvais-Jarvis, F., 2019. Emerging role of testosterone in pancreatic beta-cell function and insulin secretion. The Journal of endocrinology.

[50] Navarro, G., Allard, C., Morford, J.J., Xu, W., Liu, S., Molinas, A.J., et al., 2018. Androgen excess in pancreatic beta cells and neurons predisposes female mice to type 2 diabetes. JCI insight 3:10.1172/jci.insight.98607. eCollection 2018 Jun 21.

[51] Caldwell, A.S., Middleton, L.J., Jimenez, M., Desai, R., McMahon, A.C., Allan, C.M., et al., 2014. Characterization of reproductive, metabolic, and endocrine features of polycystic ovary syndrome in female hyperandrogenic mouse models. Endocrinology 155:3146–3159.

[52] Dunaif, A., Segal, K.R., Shelley, D.R., Green, G., Dobrjansky, A., Licholai, T., 1992. Evidence for distinctive and intrinsic defects in insulin action in polycystic ovary syndrome. Diabetes 41:1257–1266.

[53] van Schaftingen, E., Gerin, I., 2002. The glucose-6-phosphatase system. The Biochemical journal 362:513–532.

[54] Taniguchi, C.M., Emanuelli, B., Kahn, C.R., 2006. Critical nodes in signalling pathways: insights into insulin action. Nature reviews.Molecular cell biology 7:85–96.

[55] Kobayashi, A., Azuma, K., Ikeda, K., Inoue, S., 2020. Mechanisms Underlying the Regulation of Mitochondrial Respiratory Chain Complexes by Nuclear Steroid Receptors. International journal of molecular sciences 21:10.3390/ijms21186683.

[56] Levin, E.R., Hammes, S.R., 2016. Nuclear receptors outside the nucleus: extranuclear signalling by steroid receptors. Nature reviews.Molecular cell biology 17:783–797.

[57] Arkun, Y., 2016. Dynamic Modeling and Analysis of the Cross-Talk between Insulin/AKT and MAPK/ERK Signaling Pathways. PloS one 11:e0149684.

