## Supplemental data for "Hepatocyte androgen receptor in females mediates androgen-induced hepatocellular glucose mishandling and systemic insulin resistance"

**Suppl. Figure. 1**

AR Protein Expression

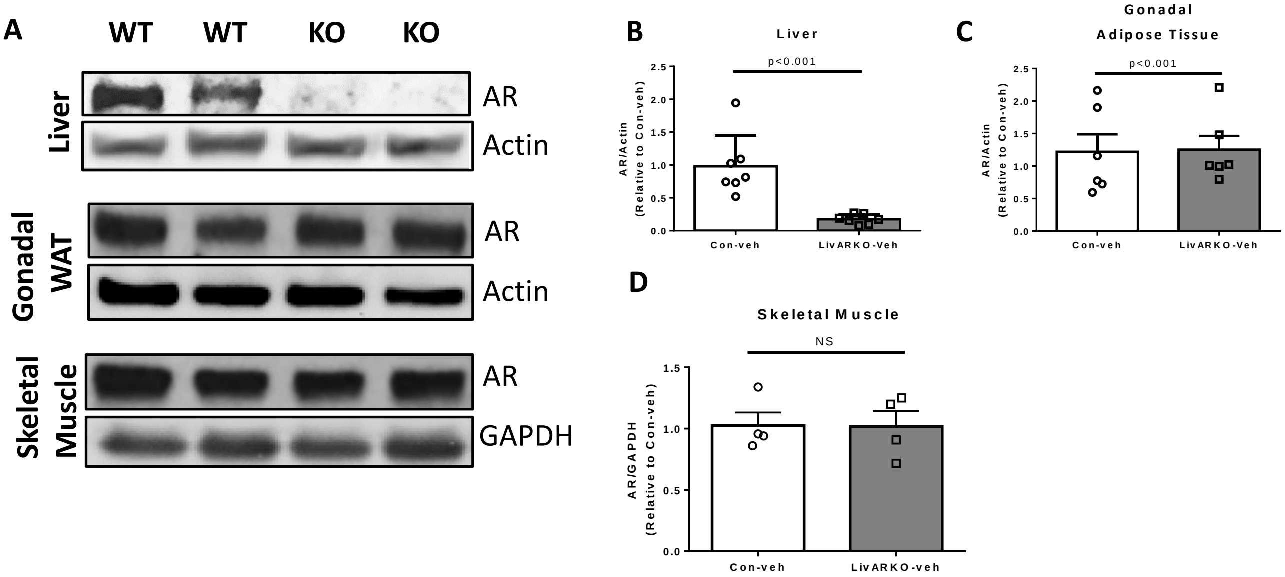


**Suppl. Figure 1. AR protein levels are significantly reduced in liver.** Western blot of AR protein (A) in liver, adipose and muscle. AR protein levels were quantified by densitometry from western blots. The AR protein levels were significantly reduced in the liver of LivARKO compared with control mice (B) without change in gonadal adipose tissues (C) or muscle (D). Two-tailed student's t-test was applied. Values are mean±S.E.M. N=4-7 per group.

**Suppl. Figure 2**

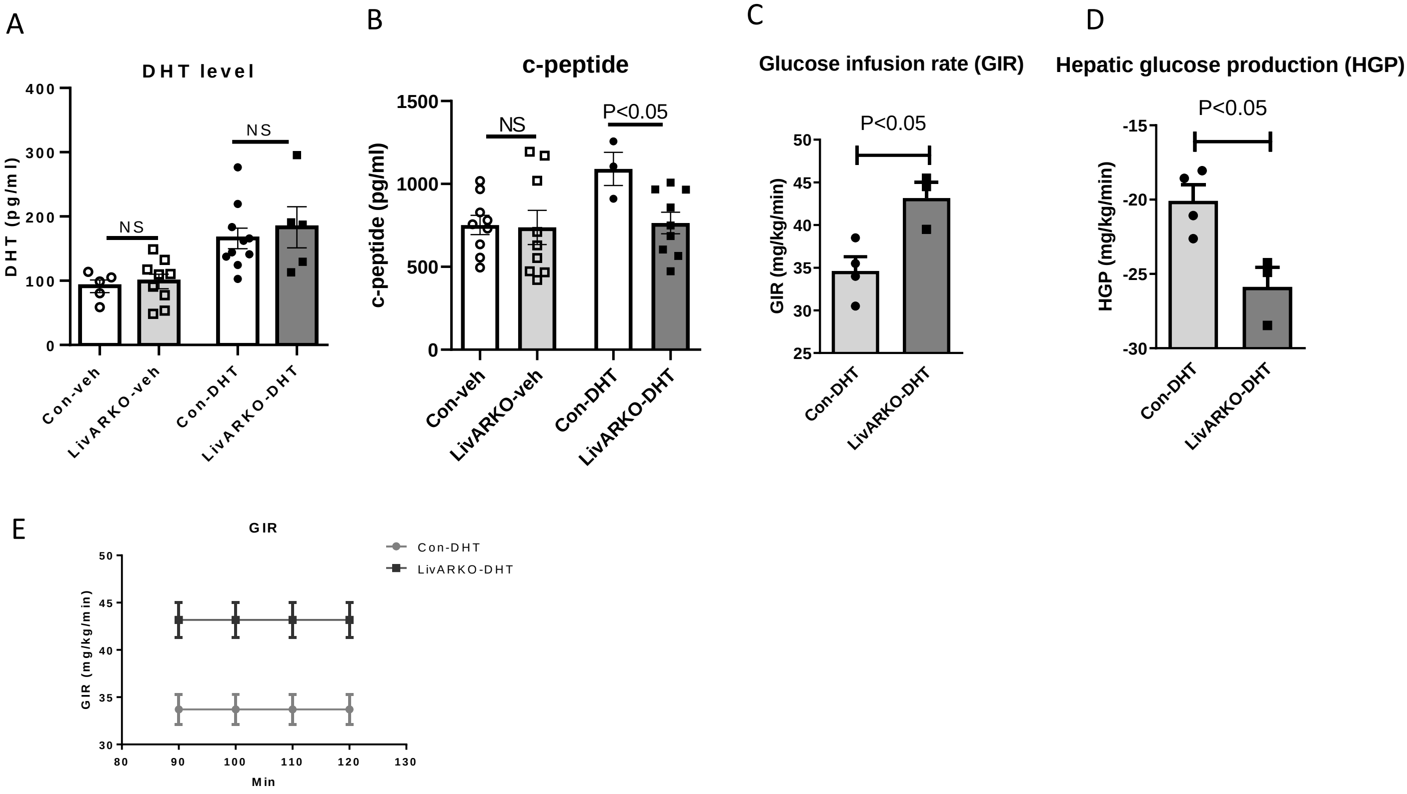


**Suppl. Figure 2.** Blood hormone levels and glucose infusion rate. Blood was collected from control and LivARKO mice with or without DHT 2-3 months postinsertion. (A) DHT levels. (B) C-peptide levels. (C) the glucose infusion rate (GIR), (D) hepatic glucose production and (E) GIR at 90, 100, 110 and 120min after insulin infusion was determined in a euglycemic-hyperinsulinemic clamp system with blood glucose levels at the steady state. Data were compared by two-tailed student's t-test. Values are means ±S.E.M.

**Suppl. Figure 3**

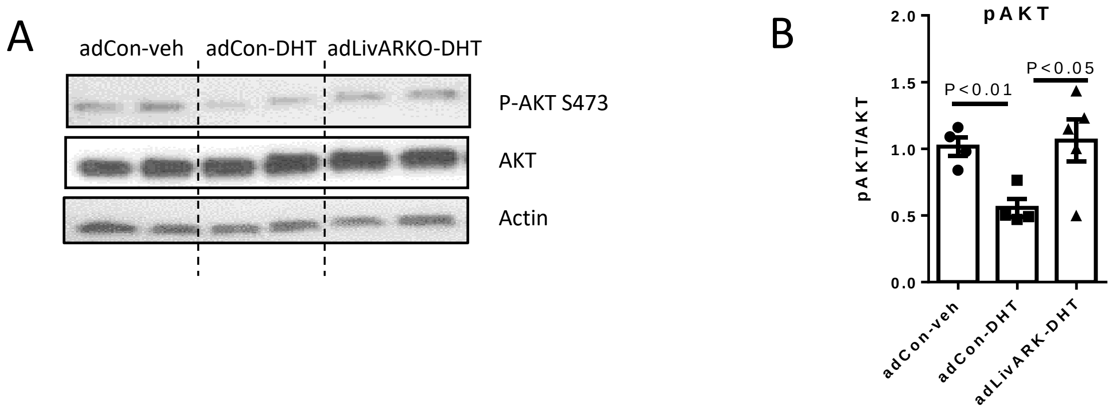


**Suppl. Figure. 3 Deletion of liver AR by AAV-Cre prevented DHT-induced insulin resistance.** At 3 months postinsertion, GFP and adLivARKO mice were fasted for 16 hours and then injected with 0.5 U/kg insulin. After 10 minutes, liver samples were collected and subjected to (A) western blot analysis and (B) The densitometry graphical representations for p-AKT (S473)/AKT. n = 3-4 per group. Data were assessed by compared by two-tailed student's t-test compared to adCon-DHT.

**Supplemental Figure 4.**

### Human Primary Hepatocytes (sample 1)

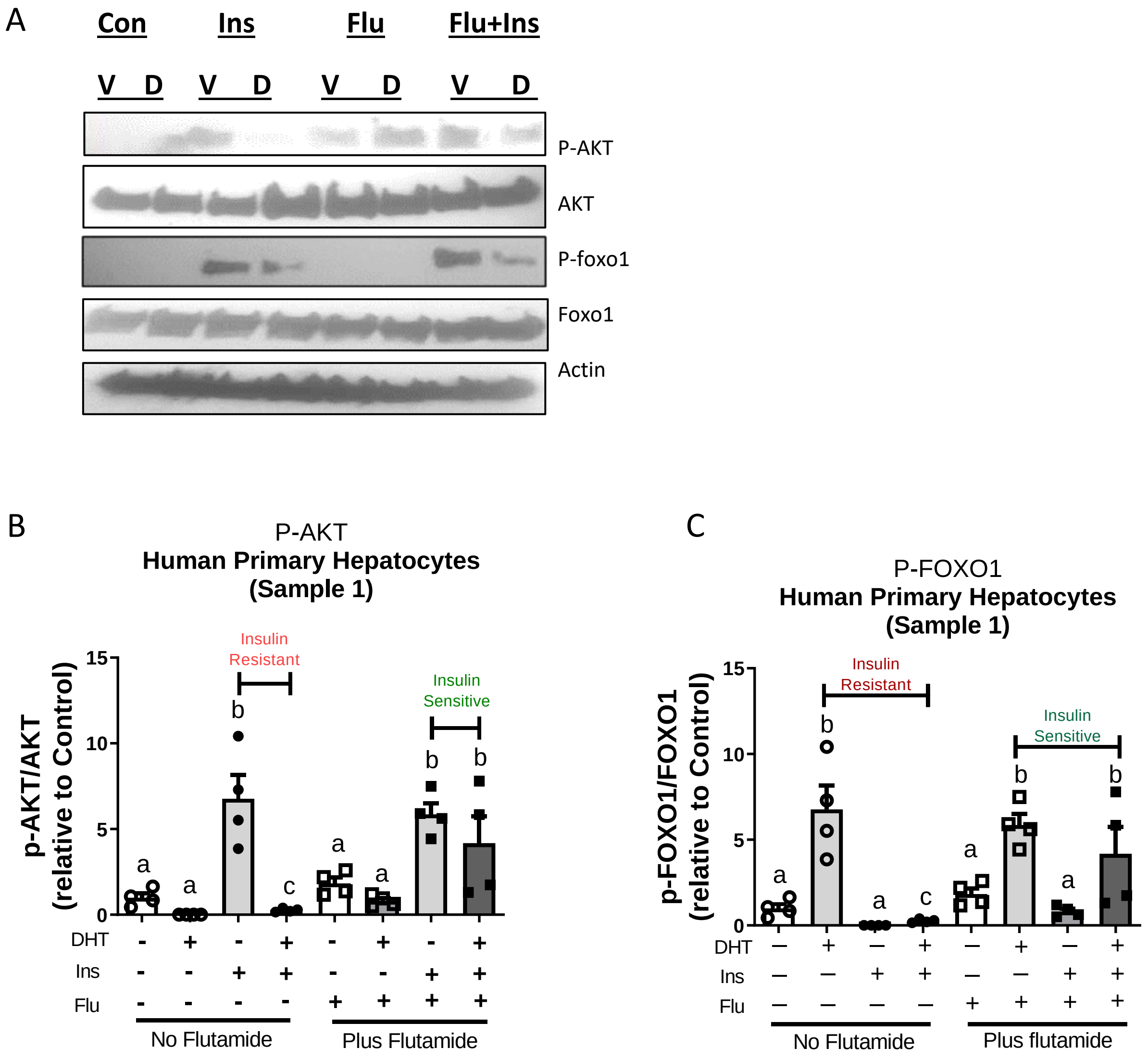

### Human Primary Hepatocytes (sample 2)

D

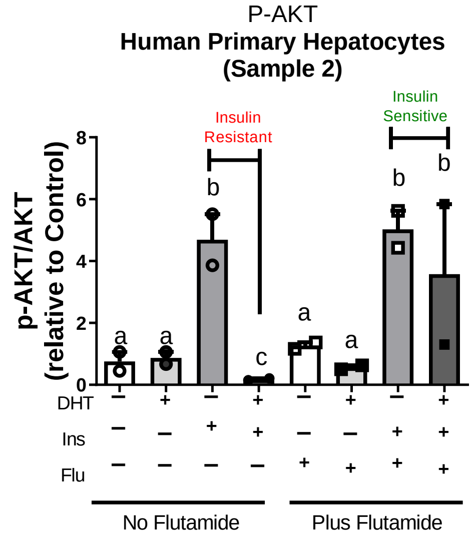


E

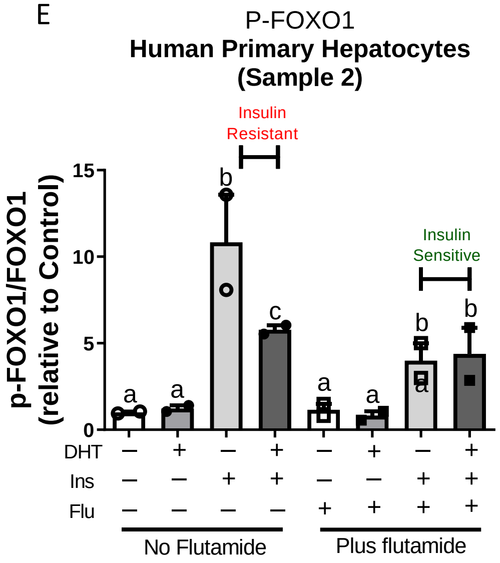


### Human Primary Hepatocytes (sample 3)

F

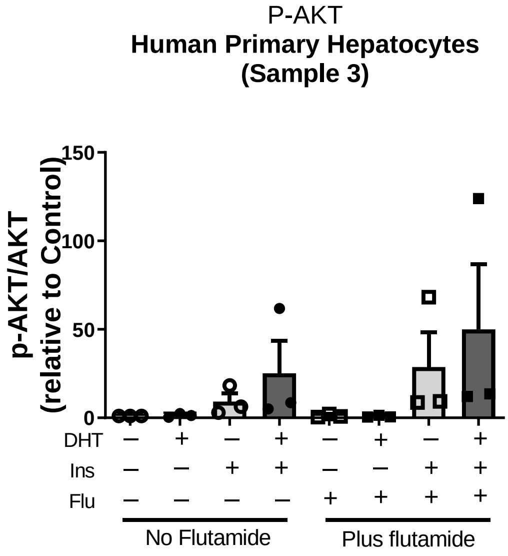


G

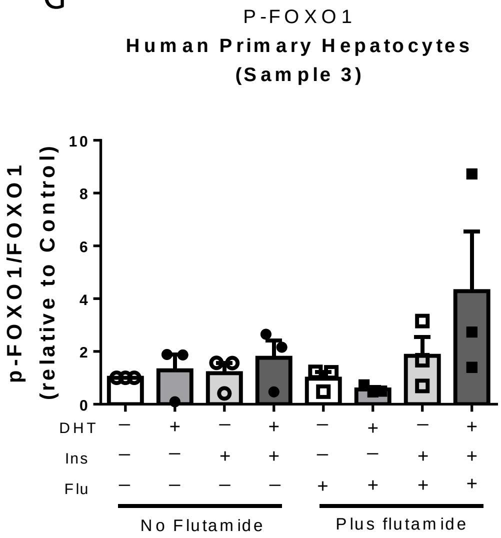


**Supplemental Figure 4** Prevention of DHT-induced impairment of hepatic insulin signaling was recapitulated in primary hepatocytes from two women.
Primary hepatocytes were extracted from three women (A-C for sample 1; D-E for sample 2; F-G for sample 3), cultured for 24 hours, subjected to a cell culture DHT treatment plus and minus flutamide, and assessed via Western blot analysis using antibodies against p-AKT, AKT, p-Foxo1, Foxo1, and Actin. Graphical representations of the densitometry for (B, D, F) p-AKT/AKT and (C, E, G) p-Foxo1/Foxo1 are shown. Each sample with 2-4 replicates. Data were compared by two-way ANOVA followed by Tukey's test. Values are mean±S.E.M. Different letter represents statistical difference, $P<0.05$.

### Suppl. Table 1 Patient information

| University of Maryland Brain and Tissue Bank | | | | | | | | | | | |
|---|---|---|---|---|---|---|---|---|---|---|---|
| | | | | Pathologist notes | | | | | | | |
| Female # | Age | Race | BMI | HIV | Hepatitis B/C | CMV | Cause of Death | Tobacco Use | Alcohol Use | Substance Use | Medical History |
| 1 | 59 | Caucasian | 24.1 | Negative | Negative | Positive | ICH | Yes | Yes | No | HTN, PAD |
| 2 | 39 | Asian | 22.9 | Negative | Negative | Positive | CNS tumor | No | No | No | CNS tumor; Dexamethasone, Keppra |
| 3 | 53 | African-American | 19.5 | Negative | Negative | Positive | CVA | Yes | Yes | No | HTN |
